# Degree bias predicts, outperforms and improves machine learning phage-host interaction predictions

**DOI:** 10.64898/2026.10.29.762857

**Authors:** Gustavo S. Jeuken, Yuval Mulla

## Abstract

Phages are promising therapeutics for difficult-to-treat bacterial infections. However, phages are highly specific in which bacterial strains they infect, necessitating extensive experimental screening for each patient isolate. Machine learning (ML) models predicting phage-host interactions from genomes are increasingly developed to prioritize phages for experimental screening. The Area Under the Receiver Operating Characteristic (AUROC) metric is standard for model evaluation, with reported high scores (0.7-0.95) suggesting strong predictive power. However, because phage-bacterium networks are highly structured, these scores can be inflated by “degree bias”, the baseline likelihood of infection driven purely by average phage infectivity and bacterial susceptibility. Here, we show that an interaction-naive model based solely on network degrees accurately predicts and outperforms the global AUROC scores of five published ML models across ten datasets, indicating that global AUROC primarily reflects network structure rather than predicted interactions. Using randomized degree-conserved networks and degree-stratified evaluation, we isolate true interaction predictions and demonstrate that current ML model performance is far more modest than global AUROC suggests. Finally, we demonstrate that regressing out degree bias from training data improves interaction predictability and boosts global and stratified performance. Our work provides a critical framework for model evaluation and a straightforward route to improve ML-guided phage selection.

## Introduction

Phages are highly specific in which bacteria they infect^1^. This specificity complicates phage therapy treatment as the phage needs to match the patient’s infecting strain^2^. This matching requiring extensive experimental testing and complex logistics when multiple biobanks are involved^3^. Recently, several machine learning models have been developed to predict infectivity from bacterial and phage genomes^4–10^. Such predictive models help prioritize which phages need to be applied and/or tested against a sequenced patient’s isolate. As the *in silico* screen does not rely on physical tests, sequenced phages from other biobanks or even *de novo* generated genomes^11^ can be considered by these machine learning models, therefore vastly expanding treatment options^12,13^.

Although model architectures vary, there is a generic approach. A phage and bacterial biobank are established, and both biobanks are fully sequenced. Next, the infectivity of all pairwise combinations of phages and bacteria is measured. Then, features are extracted from the bacterial and phage genomes. A machine learning model is trained by correlating genetic features with phage-host infectivity to train the model. After training, the model predicts how likely a sequenced phage infection is for any sequenced bacterial strain. Typically, a score is given between 0 and 1 for each in silico screened phage against the bacterial strain of interest. The highest scoring phages can subsequently be prioritized for experimental verification and potential administration to a patient.

Several metrics exist to evaluate the success of these machine learning models, with the Area Under the Receiver Operating Characteristic (AUROC) score used most prominently in the field^4–10^. To calculate this score, a model is first trained on a subset of the data. Next, the model is tested on the part of the data not used in training. The AUROC score measures a binary classification model’s ability to distinguish between positive and negative classes in the test data across all possible decision thresholds. It is calculated by plotting the True Positive Rate against the False Positive Rate (Fig. 1a). An AUROC of 1.0 indicates a model with perfect discrimination, while 0.5 represents performance no better than random guessing.

**Fig. 1.**
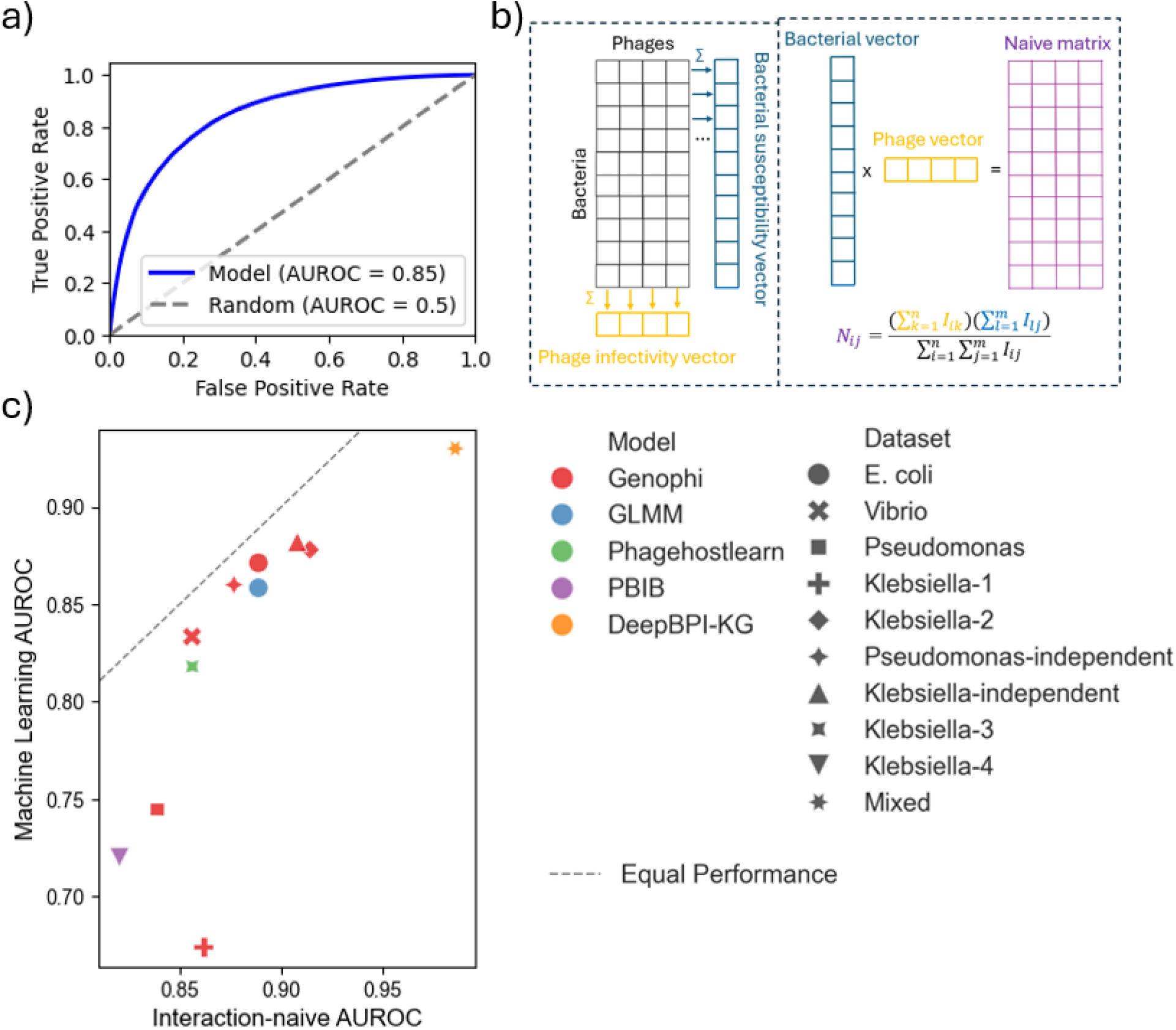
Interaction-naive model outperforms and predicts the performance of all machine learning models. **a)** Exemplary dataset (Genophi predictions^8^ on the *E. coli* dataset) showing how model performance is evaluated. The true positivity rate is integrated over the fale positivity rate to obtain the Area Under the Receiver Operating Characteristic curve (AUROC) values. Models are more predictive than random if they achieve higher AUROC scores than 0.5. **b)** From the interaction matrix (black), we construct an interaction-naive model (purple) that only considers the phage (yellow) and bacterial (blue) degree. For each phage-bacterium pair, an infectivity score is predicted based on the average bacterial susceptibility and the average phage infectivity. **c)** The naive model has a higher AUROC score than all machine learning models on all tested datasets. Furthermore, the naive AUROC is highly predictive of each model’s AUROC rank (Spearman’s r=0.89, p=10^-4^), despite having no knowledge about the models.

All current machine learning models report AUROC scores between 0.7-0.95^4–10^, suggesting moderate to high predictability. However, all phage-bacterium interaction networks are known to be highly structured^14–16^. Phages with broad host ranges in the training data (generalists) are naturally more likely to infect unseen bacteria than narrow-host-range phages (specialists), a trend that requires no machine learning model to anticipate. In predictive modeling, the performance boost driven purely by average phage infectivity and bacterial susceptibility is known as degree bias. Ultimately, a model’s reported predictive power is a combination of this structural degree bias and its actual capacity to uncover true biological interactions.

To disentangle interaction prediction from degree bias, we have compared the performance of an interaction-naive model (degree bias only) with the performance of five machine learning models^4–8^ across ten datasets^4–7,9,10,17–20^. We find that the naive model achieves higher AUROC scores than all machine learning models across all tested datasets^4–10^. Furthermore, we find that the naive model accurately predicts the ranking of all models, suggesting the AUROC score predominantly reflects network structure rather than model performance. By comparing model performance between real and degree-conserved randomized networks, we find that AUROC values are dominated by degrees and via degree-stratified evaluation we find a true but modest performance of interaction predictions beyond degrees. Lastly, we show that regressing out degree bias from the input data improves interaction predictions.

Our work highlights the importance of proper evaluation metrics, shows that current machine learning models perform more modestly in predicting phage-host interactions than perhaps previously suggested, and provides a universal route for improving ML-models in predicting phage-host interactions. Such improved models can in the future aid selection of therapeutic phages and reveal genetic determinants of phage-host interactions.

## Results

To quantify the contribution of degree bias in the performance of current machine learning models in predicting phage-host interactions, we first define an interaction-naive model that consists only of degrees (Chung-Lu configuration model^21^, consisting only of average phage infectivity and average bacterial susceptibility, Eq. 1, Fig. 1a). This interaction-naive model is not a true competitor of machine learning models as it does not do test-train splitting, but it shows for each dataset how ‘well’ a model can perform without actually predicting interactions.

We compare the performance of our interaction-naive model with that of five published machine learning models^4–8^. First, we calculate AUROC scores of the interaction-naive model across ten published phage-bacterium infectivity datasets^4–7,9,10,17–20^. Next we either calculate ourselves or take from literature the AUROC scores of five machine learning model predictions^4–10^ for these datasets. Most models only have predictions for a single dataset as they rely on species-specific knowledge in selecting features from the bacterial genomes^4–6^. Only one model (Genophi) makes predictions across multiple datasets^8–10^. We find that the naive model achieves higher AUROC scores than all models for each tested dataset (Fig. 1b). Even more strikingly, we find that the naive model AUROC score rank significantly correlates with that of the machine learning models both within (Spearman’s *r* = 0.857, *p* = 0.014, Genophi datapoints only) and between models (Spearman’s *r* = 0.875, *p* = 2 * 10^−4^, all datapoints). across all datasets, these results suggest that the AUROC score mainly reflect properties of the data (e.g. degree bias), not of the machine learning model.

We next wondered to what extent the AUROC score reflects degree bias, and whether these models predict interactions at all. We therefore not only evaluate the model predictions against the true infectivity matrix, but also against a matrix where interactions are randomized such that the degree is conserved (methods)^22^. A model that only predicts degrees will score the same on the true and randomized interactions matrix, whereas a model that truly predicts interactions will achieve a lower score on the randomized matrix.

Evaluations against randomized matrices require the model’s raw predictions for each phage-bacterium pair, which have not been published for any of the models^4–10^. We we therefore calculate the predictions of each phage-bacterium pair using the most widely applied model (Genophi)^8–10^ on a commonly used dataset in literature (*E. coli*)^17^. As a positive control, we find that the naive model’s performance does not drop upon randomization of the test data (AUROC=0.89 in both cases, Fig. 2a). By contrast, the machine learning model the learning model does drop (from 0.87 to 0.82, Fig. 2a), but maintains a high AUROC score despite randomization – further confirming degree bias dominates the AUROC score.

**Fig. 2.**
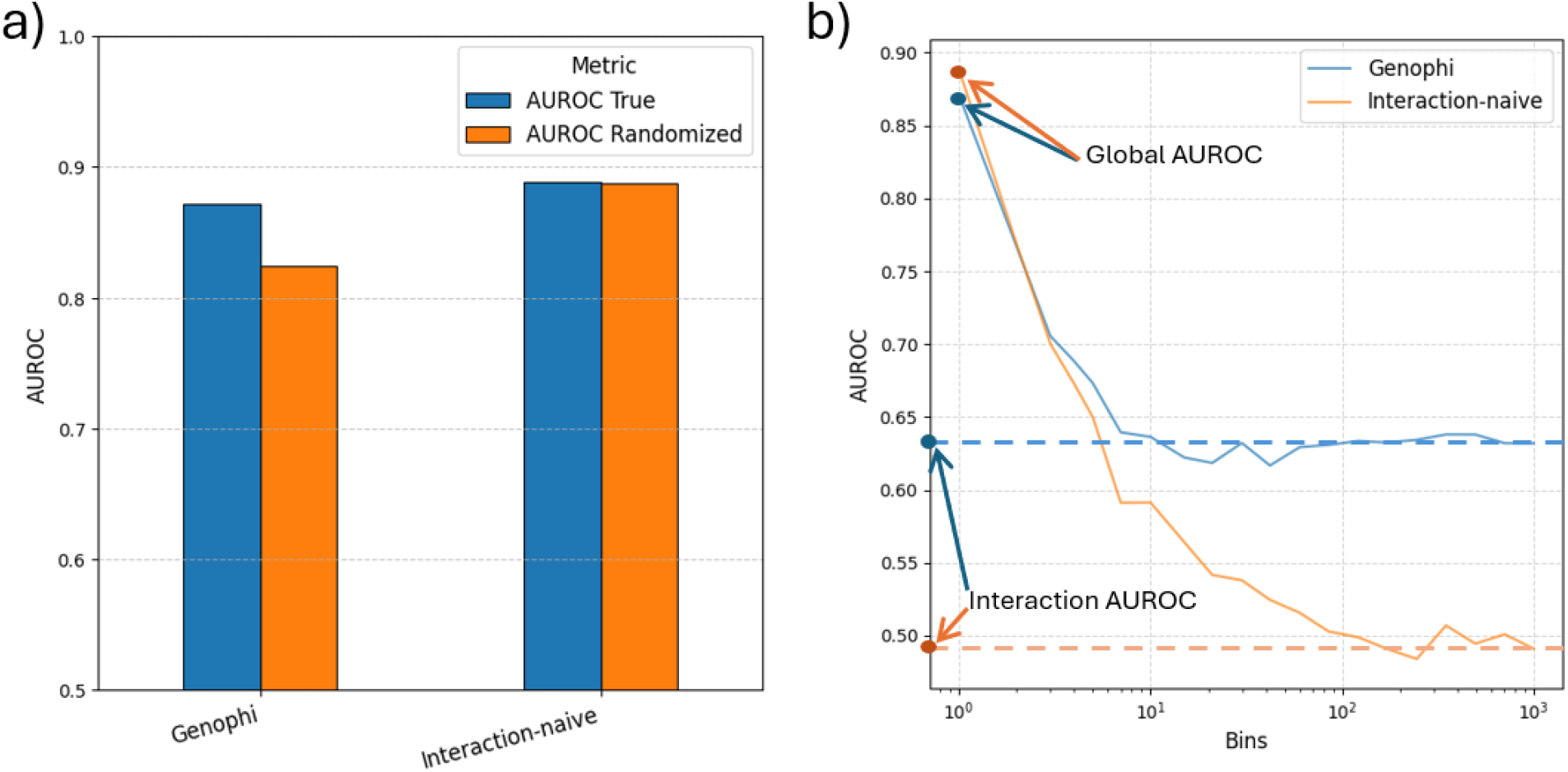
Global AUROC score is dominated by degree bias, and interaction predictability is modest. **a)** To dissect the contribution of degree bias and of true interaction predictability within the AUROC score, we create many permuted phage-host networks. For each permuted network, the interactions are randomized such that the degree bias is conserved (e.g. for each true infection taken away from a phage, a random false infection is introduced for the same phage). If a model truly predicts interactions, its AUROC score is lower on the randomized matrices than on the true matrix, whereas it stays constant if the model only. We created many randomized, degree-conserved matrices for the *E. coli* dataset^4^ and evaluated the performance of the Genophi model^8^ and of the interaction-naive model. As expected, the naive matrix model similarly between the randomized and real datasets. The machine learning model drops in performance for the randomized matrix but stays above 0.5, showing it predicts both degrees and interactions. **b**) To isolate the contribution of interactions from the global AUROC score, we perform degree-stratified evaluation by binning the AUROC score per degree. Having a single bin equals the global AUROC, whereas a highly binned evaluation isolates true interaction prediction from degree bias. Whereas the interaction-naive model globally outperforms the Genophi machine learning model, degree-stratification reveals that only the Genophi model predicts interactions – although the interaction AUROC is substantially lower than the global AUROC.

Randomization shows how well the model predicts degrees, but does not quantify how well the model predicts interactions. Therefore, we stratify the test results by degree (Methods) such that the AUROC score is exclusively reflective of the model’s capability of reflections. Whereas a bin size of 1 equals the global AUROC score from Fig. 1, increasing the number of bins isolates the contribution of interactions from degree bias. As a negative control, we evaluate our interaction-naive model and indeed find it performs comparably to random (AUROC 0.49). The machine learning model does outperform the random baseline of 0.5 (AUROC 0.63), but well below the global AUROC score (0.87). Degree-stratified performance evaluation therefore shows that the capability of the model to predict interactions is much more modest than the global AUROC score suggests.

Degree-bias is not just a problem for model evaluation; it also dominates the training objectives. We therefore test whether regressing out degree bias from the training data improves the model (Methods). Whereas interactions are binary, residuals are continuous. AUROC scores cannot be directly used to evaluate regressions. Therefore, we analyze the predictability of residuals via Pearson analysis and find a moderate correlation between real and predicted residuals of 0.388 (p<10^-10^).

To directly compare the residual-regressed with interaction-trained model, we transform the predicted residuals to interactions by using the phage degree from the training daa and predicted bacterial degrees (Methods Eq. 2). We next calculate global and degree-stratified AUROC scores for all models. We find that a machine learning model trained on degree-regressed data outperforms both a ‘normal’ model that was trained on interactions and (for the first time) the interaction-naive model (0.91 vs. resp. 0.87 and 0.89). More importantly, we find that the degree-regressed model improves the Genophi model compared to training with interactions (resp. 0.63 vs. 0.68). These result shows that regressing out degree bias from training data can improve a model’s performance in predicting phage-bacterium interactions.

## Discussion

Machine learning models hold great promise in prioritizing which phages need to be applied and/or tested against a sequenced patient’s isolate. The global AUROC score is currently the most dominantly used metric to evaluate such models^4–10^. We show that this metric mainly strongly correlates with degree bias (Fig. 1), and is poorly reflective of how well the models predict interactions (Fig. 2). Capturing degrees does not help in prioritizing phages beyond what can be achieved without machine learning. We next showed that degree-stratified evaluation isolates interaction prediction from degree bias and is therefore a useful metric in evaluating models (Fig. 2). By applying the analysis to the most used model^8^ on a commonly used dataset^4^, we show that current machine models can achieve a high global AUROC scores despite low predictability of interactions. Lastly, we show that regressing out degree bias from the training data improves model performance, providing a route towards more applicable models. We speculate that regressing out higher-order network structures from the training data can further improve models.

We used the AUROC score in different forms (global, degree-stratified, tested on a randomized matrix) to evaluate the performance of (naive and machine-learning) models because it is the most dominantly used in the field^4–10^. Other metrics such as F1, precision, accuracy recall, Matthew Correlation Coefficient (MCC) etc. are sometimes calculated as well^6–10^ but rarely reported as prominently (e.g. in the abstract) as the AUROC score. Generally, for class-imbalanced datasets such as phage-bacterium interaction matrices, where the majority does not infect, the MCC is more readily interpretable and should perhaps replace AUROC^23^. Regardless, all these metrics suffer from degree bias when calculated globally and our analysis pipeline of randomized matrices and degree-stratified testing can be directly applied to any of these evaluation metrics.

We started off with a meta-analysis of several published models tested on many experimental datasets (Fig. 1) and a more detailed analysis on the most widely used model (Genophi^8–10^) tested on a commonly used published experimental dataset (*E. coli*)^4^. Although a more wide-ranging detailed comparison was prohibited due the absence of published per-phage/bacterium predictions, it will be interesting in future work to dissect to which extent models differ in their capabilities to predict interactions– especially when tested on the same dataset^4,5,8^. Similarly, it will be interesting to see the improvement in predictions across different models when regressing out degree bias from the input data.

Despite the observed improvement (Fig. 3), machine learning model performance in predicting interactions remains moderate. We speculate that the main bottleneck for further improving machine learning predictions is not in better models, but in more training data. The currently largest dataset consists of several tens of thousands of phage-bacterium pairs. However, these networks inherently face strong class imbalance as most phages do not infect most bacteria. Furthermore, the problem of predicting infectivity from genomes is challenging as there are many bacterial genetic determinants, and because phages are highly diverse. All currently available large datasets have been captured through manual phage-host spotting^4–7,9,10,17–20^, which are incredibly laborious to perform at scale. We therefore believe that the field needs to leverage advances in robotics^24–26^, microfluidics^27–29^ and/or sequencing-derived infection networks^30–32^ to create much larger training datasets and fully unlock the power of machine learning models.

**Fig. 3.**
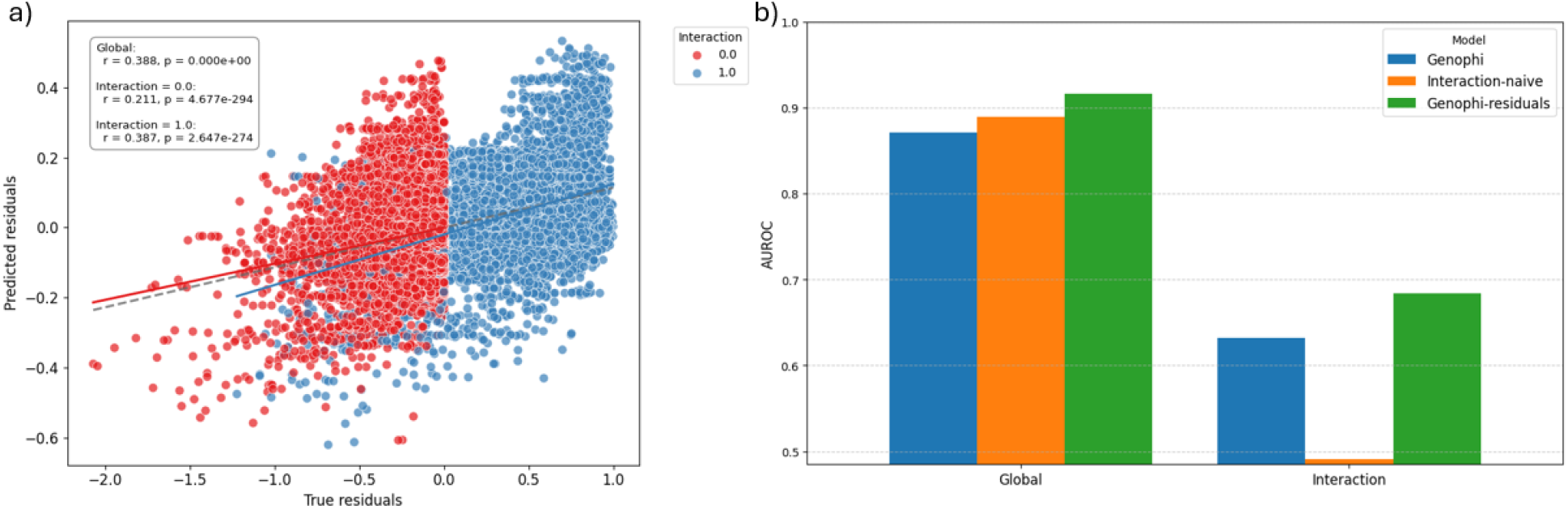
Regressing degree-bias out of the input data improves a machine learning model’s predictability of interactions. We regress out the degree bias (Eq. 2) by subtracting the interaction-naive model’s predictions from the true interactions and feed these residuals to a machine learning model. **a)** A degree-regressed Genophi model predicts both the residuals of the interacting and non-interacting phage-bacterium pairs with modest accuracy (Pearson *r* of resp. 0.387 and 0.211). **b)** We transform predicted residuals to predicted interactions by Eq. 3 to enable AUROC calculations and direct comparison with a Genophi model trained with interactions. We find that the residuals-trained model outperforms the interaction-trained model both when evaluated globally and when evaluated on interaction predictions.

The impact of such fully powered machine learning models will accelerate both therapeutic and scientific discovery. Phage therapy centers will be able to exploit (much) larger or even decentralized phage biobanks as models will help prioritize screening. Scientifically, advances in explainable AI (XAI)^33–35^ will not only help identify which phages are infective but also provide hypotheses for which genetic elements govern infectivity. The Genophi model already provides SHAP values to identify predictive features^8–10^. Following the success of genome-wide association studies (GWAS) in humans^36^, large phage-bacterium datasets combined with XAI will uncover phage biology at an unprecedented scale.

## Methods

### Construction of the interaction-naive model

We start from a 2D, binary interaction matrix *I*, where the columns present the phages and the rows present the bacteria. We first construct vector with the average phage infectivity 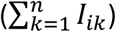 and a vector with the average bacterial susceptibility 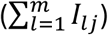. We multiply the phage and bacterial degree vectors to construct an interaction-naive matrix *N* that consists solely of expected degrees and normalize by the sum of all interactions 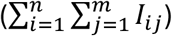 to ensure that the interaction-naive matrix is marginally calibrated:

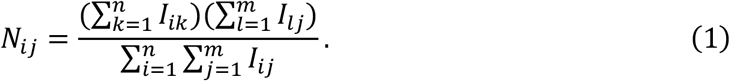

This model is the expected adjacency matrix of the bipartite Chung-Lu configuration model^21^, and we refer to it as the interaction-naïve model as the model is unaware of any specific interactions between phages and bacteria beyond their average infectivity and average susceptibility.

### Evaluation of model’s degree predictions

To isolate the contribution of degree-bias in the prediction evaluation of the model, we take the model trained on normal (non-randomized) interactions and randomized the interactions in the test data, keeping the degrees fixed. The AUROC score achieved on this randomized dataset exclusively reflects the ability of a model to capture degrees, and does not include any interaction predictions. To perform the randomization, we use previously published ‘curveball’ algorithm^22^.

### Evaluation of model’s interaction predictions

To isolate the contribution of predicting degrees from degree-bias in the evaluation of the model, we stratify the test data according to the product of the bacterial and phage degree (i.e. the predicted value for each bacterium-phage pair by the interaction-naive model, Eq. 1). Across each bin, we calculate the AUROC value and we average all AUROC scores proportionally to the number of samples per bin.

Stratification inherently involves a finite bin size. When the bin size is too small, the calculated AUROC values become very noisy due to limited number of samples er bin, whereas too large bin sizes maintain substantial degree variation within the bin. We used a bin size of 1/30 (30 equally spaced bins) as balance. Indeed for this bin size we observe an AUROC around 0.5 for the negative control of the interaction-naïve model and we observe a relatively constant AUROC score across bin sizes for the model predictions, which is only achievable if the bin size is large enough.

### Regressing degree bias out of the input data

We regress out the degree bias from the training data by subtracting the naIve matrix N (Eq. 1) interaction matrix I:

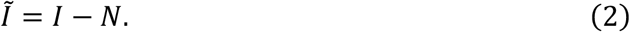

Next, we train the model (Genophi) using regression instead of classification as 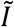 is continuous as opposed to binary. As the model is trained on residuals rather than interactions, it also predicts residuals rather than interactions. To compare the model performance across the same dataset (*I*) we therefore need to add the naive expectation *N* to the predicted residuals. For the phage degree and the average infectivity across the matrix, we can use the values from the training data. However, the model is evaluated using unseen bacteria as this mimics the condition of a new patient isolate for which a previously tested phage bank needs to be evaluated. Therefore, the bacterial degree is unknown for the entire test dataset. To solve this ‘cold start’ problem, we separately use Genophi to predict the average susceptibility for each bacterium by only feeding it bacterial genomes and their average susceptibility. Lastly, we use the average phage infectivity and average total infectivity from the input data and predicted bacterial degree to transform the residuals predictions to interaction predictions.

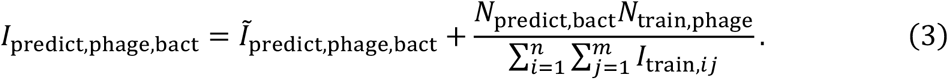

Where 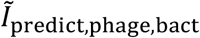 are the predicted residuals, *N*_predict,bact_ the predicted bacterial degrees, *N*_train,phage_ the phage degrees fin the training data and *I*_train,*ij*_ the interaction matrix used in the training data (before subtracting the interaction-naive matrix). We evaluate the predicted interactions (*I*_predict,phage,bact_) against the true interaction matrix *I* identically to how we evaluate predictions from the model trained with the interaction matrix directly rather than with the residuals.

